# Towards integration of time-resolved confocal microscopy of a 3D *in vitro* microfluidic platform with a hybrid multiscale model of tumor angiogenesis

**DOI:** 10.1101/2021.09.29.462293

**Authors:** Caleb M. Phillips, Ernesto A. B. F. Lima, Manasa Gadde, Angela M. Jarrett, Marissa Nichole Rylander, Thomas E. Yankeelov

## Abstract

The goal of this study is to calibrate a multiscale model of tumor angiogenesis with time-resolved data to allow for systematic testing of mathematical predictions of vascular sprouting. The multi-scale model consists of an agent-based description of tumor and endothelial cell dynamics coupled to a continuum model of vascular endothelial growth factor concentration. First, we calibrate ordinary differential equation models to time-resolved protein expression data to estimate the rates of secretion and consumption of vascular endothelial growth factor by endothelial and tumor cells, respectively. These parameters are then input into the multiscale tumor angiogenesis model, and the remaining model parameters are then calibrated to time resolved confocal microscopy images obtained within a 3D vascularized microfluidic platform. The microfluidic platform mimics a functional blood vessel with a surrounding collagen matrix seeded with inflammatory breast cancer cells, which induce tumor angiogenesis. Once the multi-scale model is fully parameterized, we forecast the spatiotemporal distribution of vascular sprouts at future time points and directly compare the predictions to experimentally measured data. We assess the ability of our model to globally recapitulate angiogenic vasculature density, resulting in an average relative calibration error of 17.7% ± 6.3% and an average prediction error of 20.2% ± 4% and 21.7% ± 3.6% using one and four calibrated parameters, respectively. We then assess the model’s ability to predict local vessel morphology (individualized vessel structure as opposed to global vascular density), initialized with the first time point and calibrated with two intermediate time points. To the best of our knowledge, this represents the first study to integrate well-controlled, experimental data into a mechanism-based, multiscale, mathematical model of angiogenic sprouting to make specific, testable predictions.

## 1 Introduction

There is now a mature literature on the mathematical modeling of tumor initiation, expansion, angiogenesis, and invasion [1–12]. These models (which may be discrete, continuous, or hybrid) seek to provide new strategies for understanding the underlying biology, and then use this knowledge to make predictions of the spatiotemporal evolution of the disease as well as its response to therapy [13–20]. A fundamental barrier limiting progress in the field, though, is the paucity of studies linking mechanism-based mathematical models with the appropriate experimental data [21–25]. One fundamental reason for this impasse is that most mathematical models require specific and quantitative data sets to calibrate model parameters; data types that are not frequently available. This is especially true in the field of tumor angiogenesis where the mathematical models demand data that are acquired at both high temporal and spatial resolution to resolve vascular dynamics [22, 26–30]. Furthermore, the models themselves may have a myriad of parameters yielding an ill-posed parameter calibration problem [31–34]. Still, these models, even if uncalibrated against experimental data, can serve as useful tools to guide the experimental study of tumor-induced angiogenesis. For example, Vilanova *et al*. developed a phase-field model of capillary progression with a discrete tip endothelial cell that guides capillaries based on tumor angiogenic factors and neighboring capillaries [35–37]. This model has the ability to account for capillary regression due to anti-angiogenic therapies and qualitatively matches *in vivo* experimental measurements. Their approach was extended by Xu *et al*. who incorporated blood flow and initialized the resulting model with photo-acoustic imaging data characterizing the tumor and surrounding vasculature [38]. Travasso *et al*. developed a phase-field approach describing endothelial, stalk, and tip cells that compares differences in vessel morphology due to tip cell migration and stalk cell proliferation [39]. Their model’s predictions were qualitatively compared to experimental data of vascular patterns in response to the production level of angiogenic factors. These efforts are to be commended as some of the first examples of linking mechanism-based, mathematical models of tumor angiogenesis with experimental data. The next step is to calibrate such a model with early time point data, and then make a prediction to directly compare model forecasts to data acquired experimentally at later time points.

Calibrating mathematical models with experimental data allows for the ability to both predict future model outcomes and more rigorously test and understand the biological phenomena in question [23, 24, 40, 41]. Specifically, for tumor angiogenesis, the ability to calibrate parameters that dictate the development of tumor vasculature in a well-controlled, experimental platform would allow the systematic testing of various models of angiogenesis [21, 22]. However, obtaining the appropriate data for this task remains a difficult challenge. A substantial focus of experimental angiogenesis research is performed *in vitro* on 2D monolayers as well as *in vivo* xenograft models in animals. However, 2D systems do not mimic the complex behaviors observed *in vivo*, while xenograft systems pose substantial barriers to acquiring the data with adequate spatial and temporal resolution to resolve tumor induced angiogenesis [22, 42]. We do note other *in vitro* works in experimental angiogenesis [43–47]. In the present effort, we address the data demands *via* a 3D *in vitro* microfluidic platform [48]. This system has the advantage of emulating the complex architecture of the tumor microenvironment observed *in vivo*, as well as capturing the spatiotemporal behavior of the cells. The 3D *in vitro* microfluidic platform we utilized in this study consists of a collagen matrix seeded with inflammatory breast cancer (IBC) cells around a functional blood vessel to represent the IBC tumor-vascular interface. Over time, the IBC cells secrete pro-angiogenic factors that diffuse and bind to the functional blood vessel, initiating angiogenic sprouting. Spatiotemporal measurements of cell proliferation and apoptosis, branch lengths and number, anastomosis, and lumen formation in the platform will provide key parameters that drive the mathematical model.

In this contribution, we aim to calibrate an agent-based model of tumor angiogenesis [49] with experimental data obtained from the *in vitro* vascularized tumor platform [48] described above to predict vasculature sprouting. First, we perform experiments that isolate key components in the process of angiogenesis and use these data to calibrate important parameters in mathematical models of increasing complexity. This sequential approach allows a systematic, stepwise, approach for linking mathematical models with experimental data across spatial scales. The steps (referred to as scenarios in the remainder of this work) we follow are: 1) calibration of tumor and endothelial cell number to time-resolved hemocytometry, 2) calibration of the secretion and consumption of vascular endothelial growth factor (VEGF) by tumor and endothelial cells, respectively, to time-resolved protein expression measurements, 3) calibration of stalk cell growth rate against time-resolved measurements of angiogenic sprout length, performing a 4) global and 5) local spatiotemporal calibration of the hybrid model of tumor angiogenesis to confocal microscopy data acquired from the in vitro microfluidic platform, and 6) testing the predictive capabilities of the calibrated mathematical model at the global and local scales.

## 2 Methods

### 2.1 Overview of Methods

Fig 1 summarizes the computational and experimental methodologies utilized in this study. In Panel (A), we show the coupled ordinary differential equation (ODE) models to describe tumor (IBC3) and endothelial (TIME) cell number, as well as the production of VEGF by tumor cells and the consumption of VEGF by endothelial cells. These models are calibrated using cell number over time (Scenario 1) counted *via* a hemocytometer, and VEGF concentration (Scenario 2) measured *via* ELISA analysis, shown in Panel (B). Panel (C) depicts the hybrid multiscale model, initialized from confocal microscopy images observed in our 3D in vitro microfluidic platform, shown in Panel (D), and calibrated by subsequent time points. In our global analysis, shown in green in Panels (C) and (D), we calculate global quantities of interest from the confocal microscopy images, after thresholding for area and intensity and projecting the data into a 2D plane. From this 2D image, we calculate the sprout length, vascular density, and vascular volume fraction, which are readily comparable to the angiogenesis model to calibrate and predict the global sprout elongation rate (Scenario 3) and the distance between tip cells for activation (Scenario 4). After analyzing the model globally, we assess the ability of the model to recapitulate local vascular features. In our local analysis, shown in peach in Panels (C) and (D), we select individual sprouts for calibration through a Dice and area threshold over time, and skeletonize the observed vessels (and observe adjacent tumor cells). These skeletonizations, along with the tumor cells, are used to initialize and calibrate the local sprout elongation rate. We then assess the ability of the angiogenesis model to locally predict the final time point (Scenario 5). Finally, we use the calibrated parameter distributions to predict sprout length, vascular density, and vascular volume fraction (globally) and the local structure of the angiogenic sprouts.

**Figure 1.**
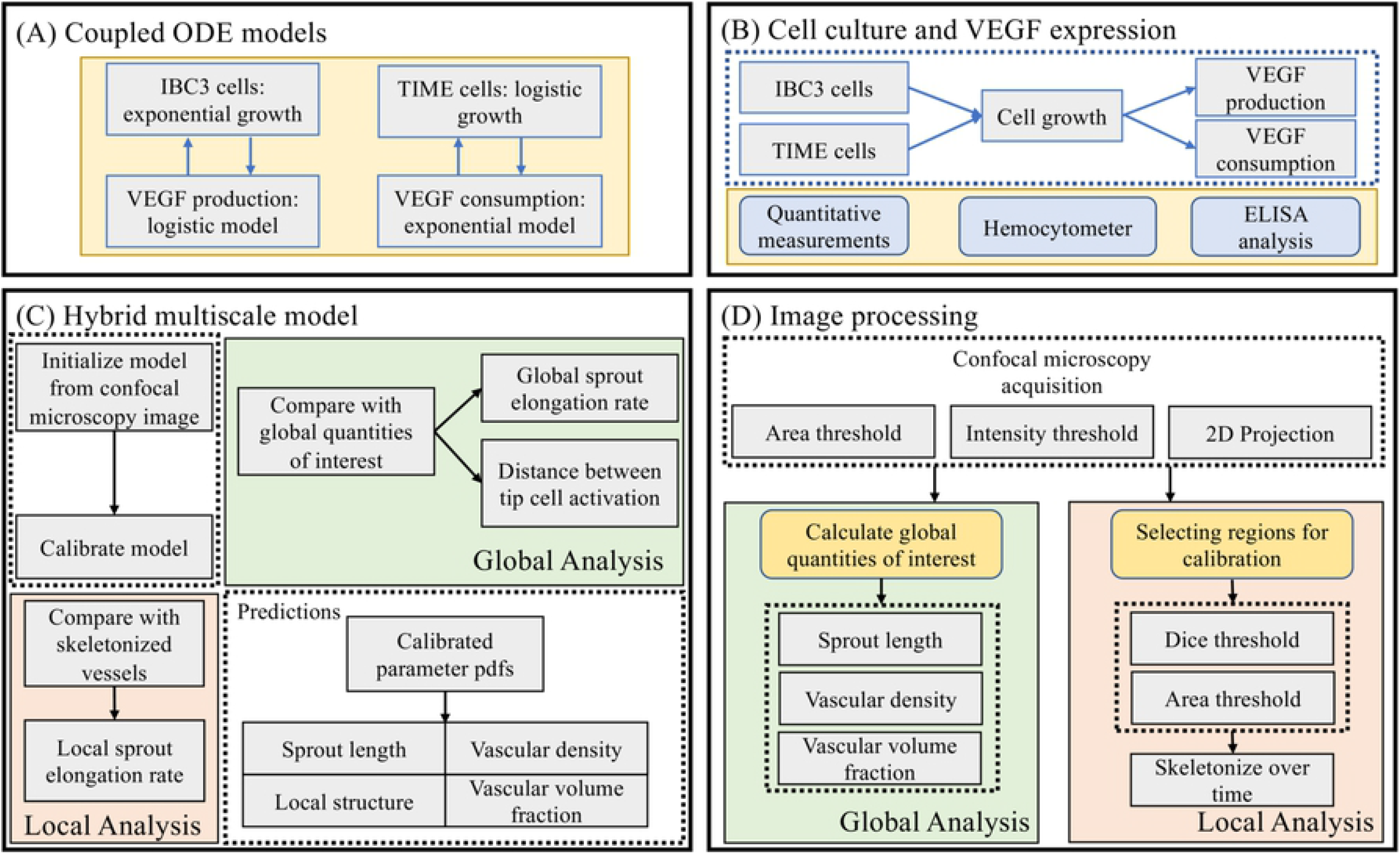
Schematic of computational and experimental methods. Panels (A) and (B) show the calibration of tumor and endothelial cell number and VEGF expression to hemocytometer and ELISA measurements over time. Given the calibrated VEGF production and consumption rates, we inform our hybrid multiscale model, shown to Panel (C), to confocal microscopy images of angiogenic sprouts, depicted in Panel (D). We analyze the model globally, calibrating model parameters to summary statistics of the data, namely the sprout length and vascular density, and locally, calibrating the local sprout elongation rate to segmented vascular structures. This sequential approach, starting with protein expression and cell number experiments to inform the hybrid model prior to integrating confocal microscopy images, allows us to utilize experimental data at multiple scales to inform the multiscale nature of the tumor angiogenesis model.

### 2.2 Experimental Methods

#### 2.2.1 Cell culture and VEGF expression

Human inflammatory breast cancer (IBC) MDA-IBC3 cells and telomerase-immortalized human microvascular endothelial (TIME) cells were used in this study. Green fluorescence protein labelled MDA-IBC3 (a HER2+ IBC cell line) were kindly provided by Dr. Wendy Woodward (MD Anderson Cancer Center, Houston, TX) and mKate labeled TIME cells were a generous gift from Dr. Shay Soker (Wake Forest Institute for Regenerative Medicine, Winston-Salem, NC). MDA-IBC3 cells were cultured in Ham’s F-12 media supplemented with 10% fetal bovine serum, 1% antibiotic-antimycotic, 1 *µ*g/ml hydrocortisone, and 5 *µ*g/ml insulin. TIME cells were cultured in Endothelial Cell Growth Medium-2 BulletKitTM (EGM-2, Lonza). All cell cultures utilized in this study were maintained at 5% CO2 atmosphere and 37°C.

To determine the production of VEGF by MDA-IBC3 cells and consumption of VEGF by TIME cells, cells were seeded with a density of 30,000 cells/cm^2^ and 10,000 cells/cm^2^, respectively, in separate 12 well plates supplemented with EGM-2 media. Media was collected and replaced from the same 4 wells daily for measurement of VEGF concentration and cell growth was imaged with an IncuCyte Zoom (Essen Bioscience, Ann Arbor, MI) every 8 hours for 7 days. VEGF concentration was measured using a Human VEGF Quantikine ELISA Kit (R&D Systems, Minneapolis, MN). The cell number in each well was determined by trypsinization and counting of cells with a hemocytometer on days 1, 2, 3, 5, and 7. For each timepoint, four wells were sacrificed for cell counting and these were separate from the media collection wells for VEGF measurements.

#### 2.2.2 Vascularized 3D *in vitro* microfluidic platforms

The *in vitro* 3D tumor microfluidic platforms utilized in this study were composed of extracellular matrix comprised of collagen type I seeded with green fluorescence protein labeled MDA-IBC3 around a hollow vessel lined with mKate labeled TIME cells housed in a polydimethylsiloxane scaffold as described in our published work. 7 mg/ml collagen solution, which has a similar stiffness to that of *in vivo* breast tumors, was used in fabricating extracellular matrix of the platforms [50–52]. MDA-IBC3 cells were seeded at a density of 1 × 10^6^ cells/ml in the collagen solution and polymerized around a 22 G needle at 37°C for 25 minutes. After polymerization, the needle was removed, and the resulting hollow vessel was filled with a solution of 2 × 10^5^ TIME cells to form an endothelialized vessel lumen. Flow was introduced using a syringe pump system and a graded flow protocol was used to establish a confluent endothelium as we have previously published [48, 51, 53–55]. The microfluidic platform is summarized in Fig 2. Panel (A) shows an example microfluidic platform with an axial cross-section depicting the functional parent blood vessel and cancer cells seeded throughout the microenvironment in Panel (C). The platforms are connected to a syringe pump system, shown in Panel (B), that allows for the continuous perfusion of media into the collagen microenvironment. Over time, the cancer cells release pro-angiogenic proteins, which diffuse through the microenvironment and cause tumor-induced angiogenesis. An example of these sprouts is shown in Panel (D).

**Figure 2.**
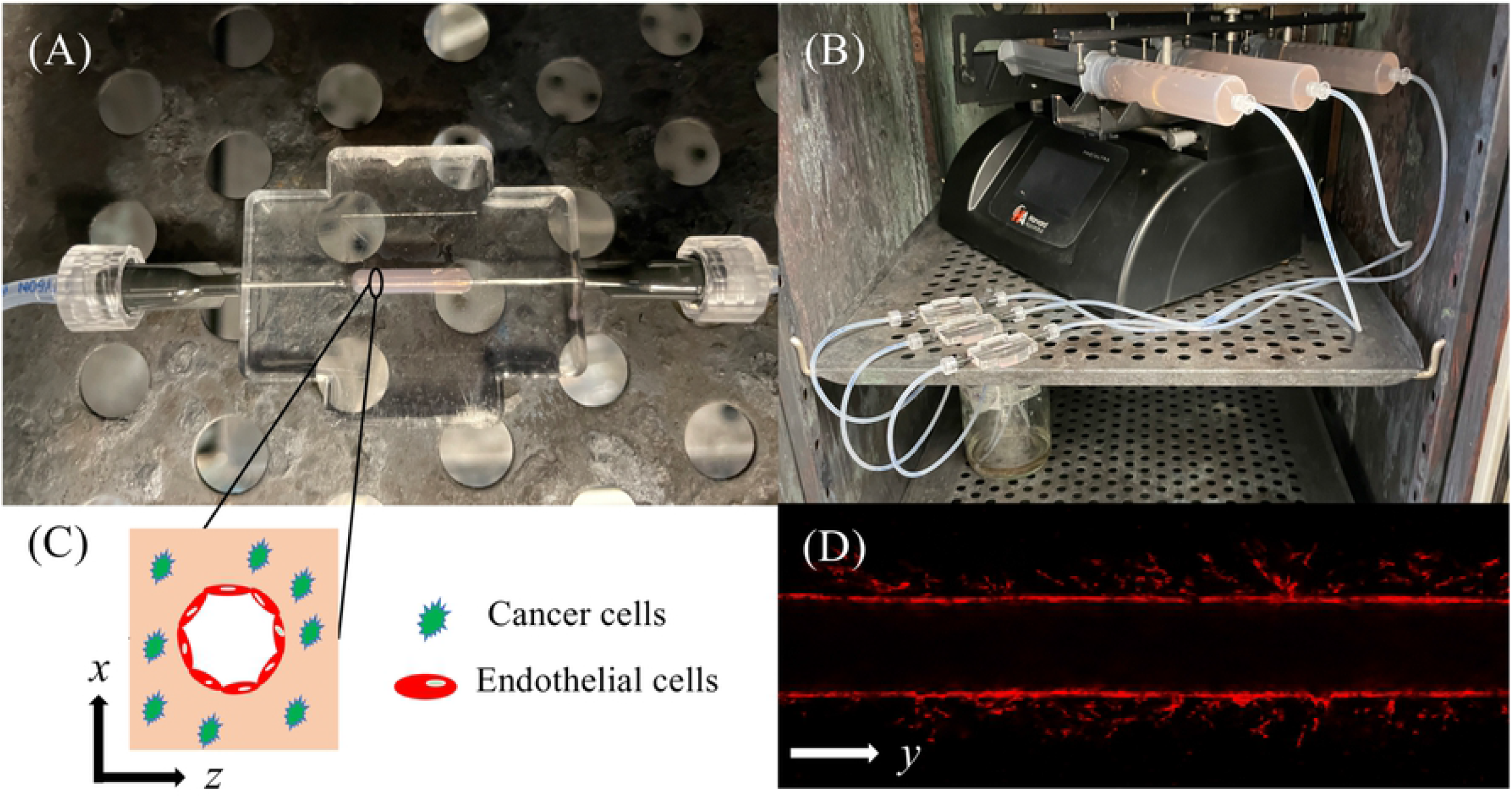
Schematic of microfluidic platform. Panel (A) depicts the housing of the 3D microfluidic platform. The black oval denotes the boundary of the platform containing collagen matrix (shown in light pink) within the housing chamber. The chamber is connected to a syringe pump flow system, shown in Panel (B), to maintain continuous flow of media through the parent vessel, within the collagen matrix. The axial cross-section of the vessel (the x-z plane) is shown in Panel (C) and an example z-slice image (i.e., longitudinal cross-section along the length of the vessel (the x-y plane)), is shown in Panel (D).

#### 2.2.3 Image acquisition and processing

Platforms were imaged with a Leica TCS SP8 Confocal Microscope (Leica Microsystems, Germany) on days 3, 5, 7, 9, 11, 15, and 19. The 3D volume of angiogenic sprouts was imaged with *z*-slices acquired at intervals of 4.28 microns and an in-plane resolution of 2.254 microns × 2.254 microns in the *x-y* plane (shown in Panel (D)). The confocal microscopy images were processed at each time point to ensure the same local structures of the microfluidic platform could be compared over the duration of the experiment. This was done *via* a reference mark. It was assumed that the diameter of the parent vessel did not deviate significantly over the course of the time points used for calibration. The parent vessel periphery was used as a reference to align the segmented region of sprouting in *x-y* space, depicted in Panel (D). We assume that the vessels grow primarily in *x-y* space and that little growth is in the *z* direction (we comment on this assumption in the Discussion section). We average signal intensity across 10 slices in the *z* direction (approximately 43 microns in total) and threshold using area and intensity (to filter out migratory endothelial cells that are not a part of maturing vessels) to create a binarized vessel mask. This mask is used to calculate both the global (i.e., sprout length, vascular density, and vascular volume fraction), and local vessel morphologies (i.e., the structure of specific vasculature) over time for local calibrations.

To select specific vessels for local calibration, we developed an algorithm that calculates the Dice and percentage of overlap across time points. This allows for selecting regions that, when aligned, correspond to angiogenic sprouts growing over time and minimizes the effects of random endothelial cell migration. For the selected regions, we utilized a Zhang-Suen thinning algorithm [56] (available at https://github.com/linbojin/Skeletonization-by-Zhang-Suen-Thinning-Algorithm) for skeletonization after dilating the vessels using the ‘strel’ function in MATLAB (Mathworks, Natick, MA). Given the skeletonization for the model and the data at time *t*, we calculate the average distance between the model to the data centerline and from the data to the model centerline to use as the objective function for model calibration by using Python’s SciPy Euclidean distance transform [57].

### 2.3 Computational methods

#### 2.3.1 Overview of computational methods

We calibrate three mathematical models for the purpose of predicting vascular response to the secretion of VEGF by tumor cells. (We first describe them qualitatively before introducing their mathematical representations.) The first and second are systems of ordinary differential equations that describe VEGF consumption by TIME cells and production of VEGF by MDA-IBC3 cells. These models describe the concentration of VEGF over time and the evolving cell type associated in production or consumption. The first model describes VEGF consumption of endothelial cells and has three model parameters: the consumption rate of VEGF by endothelial cells, and the growth rate and carrying capacity of endothelial cells. All of these parameters may be determined from the VEGF expression and cell culture data. The second model describes the growth of MDA-IBC3 cells and their production of VEGF through three parameters: the VEGF production rate and carrying capacity, and the growth rate of the tumor cells. These parameters may also be determined from the VEGF expression and cell culture data. We utilize these parameters as inputs into the third model, a hybrid multiscale model of tumor angiogenesis. An agent-based model describes the tumor and endothelial cell dynamics and a continuum model governs the dispersion of VEGF. This hybrid model predicts the spatiotemporal distribution of vascular sprouts and is readily comparable to the confocal microscopy images (and the calculated quantities of interest at the local and global scales) of the 3D *in vitro* microfluidic platform. These three models vary in complexity and allow us to isolate model parameters in a series of well-controlled experiments, and ultimately to test the predictive capabilities of the multiscale model at the global and local scales.

#### 2.3.2 Model of VEGF production and consumption

The following two systems of mathematical models describe the consumption and production of VEGF by TIME and MDA-IBC3 cells, respectively. The first system of ODEs is:

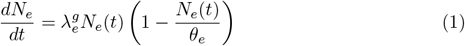

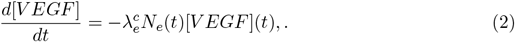

where 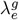 is the growth rate of endothelial cells, *N*_*e*_(*t*) is the number of endothelial cells at time *t, θ*_*e*_ is the carrying capacity of endothelial cells, [*V EGF*] is the concentration of VEGF, and 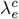 is the consumption rate of endothelial cells. This system assumes that each endothelial cell consumes VEGF at a constant rate, VEGF is not produced by endothelial cells, *N*_*e*_(*t*) follows logistic growth with carrying capacity *θ*_*e*_, and VEGF decay is negligible. To model the VEGF secretion by tumor cells we write:

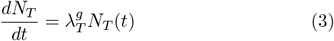

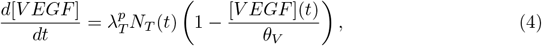

where 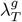 is the growth rate of tumor cells, *N*_*T*_ (*t*) is the number of tumor cells, 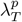 is the production rate of VEGF by tumor cells, and *θ*_*V*_ is the carrying capacity of VEGF. Since both *N*_*e*_(*t*) and *N*_*T*_ (*t*) are measured daily, we calibrate 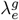 and 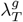 and utilize the model to predict cell numbers between experimental measurements. To directly compare model predictions to the cell culture and protein expression data, we take *N*_*T*_ (*t*) and *N*_*e*_(*t*) to be MDA-IBC3 cells and the generic endothelial cells to be TIME cells, respectively.

#### 2.3.3 Hybrid multiscale model of tumor angiogenesis

We utilize the hybrid multiscale model of tumor induced angiogenesis developed in [49] to compare the global and local quantities of interest with the confocal microscopy data in [48]. We now summarize the salient features of this model and refer the interested reader to [49]. The multiscale model describes tumor and endothelial cell dynamics using an agent-based model, and the evolution of nutrients and vascular endothelial growth factor (VEGF) using a continuous partial differential equation (PDE) model. Tumor cells act as individual agents and may reside as a proliferative, quiescent, hypoxic, or necrotic cell. They transition from quiescent to proliferative stochastically with a linear dependence on nutrient concentration. Tumor cells uptake nutrients, grow, and proliferate until the supply of nutrients is depleted (i.e., less than a threshold, *σ*_*H*_), at which point they become hypoxic and secrete VEGF. The VEGF diffuses across the domain and eventually contacts nearby vasculature. The blood vessels, comprised of endothelial cells modeled as agents, uptake VEGF and become activated when the concentration of VEGF is greater than a threshold, *α*_*A*_, and the distance between other tip cells is greater than *d*_*tip*_. The activated endothelial cells then transition to tip cells, which are responsible for the migration of angiogenic sprouts up the concentration gradient of VEGF due to a chemotactic force,

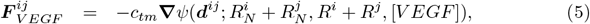

where *f*_*v*_ is the VEGF force coefficient, *d*^*ij*^ the distance between tip cell *i* and the neighboring endothelial cell *j*, 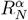 is the nuclear radius of cell *α, R*^*α*^ is the cytoplasmic radius of cell *α*, and the VEGF potential function ∇*ψ* is given as

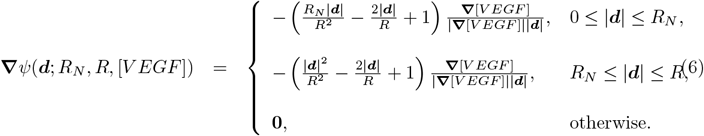

The proximal endothelial cells, termed stalk cells, proliferate at the rate *d*_*sc*_ and allow the elongation of the sprouts. Mechanical adhesion and repulsion forces maintain the structural integrity of the vessels and allow the formation of the lumen. Once the sprouts mature, they form complex networks and establish blood flow, enabling the delivery of new nutrients to the tumor. The VEGF and nutrient dynamics are governed by the following reaction-diffusion equations:

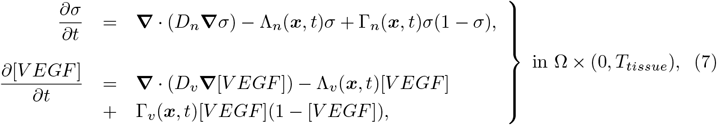

where *D*_*n*_ and *D*_*v*_ are the nutrient and VEGF diffusion coefficients respectively, Λ_*n*_ is the nutrient uptake rate of tumor cells, Λ_*v*_ is the VEGF consumption rate of endothelial cells, Γ_*n*_ is the nutrient production rate from endothelial cells that are part of anastomotic loops, Γ_*v*_ is the VEGF production rate of hypoxic tumor cells. These source and sink terms, which couple the cellular agent-based model to the continuous PDE model at the tissue scale, are defined as follows:

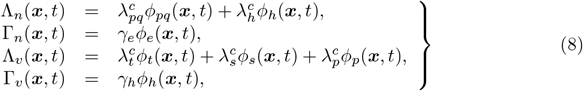

where the subscripts pq, h, l, t, s, and e specify proliferative and quiescent tumor cells, hypoxic tumor cells, looped endothelial cells, tip endothelial cells, stalk endothelial cells, and endothelial cells, respectively. Also, *ϕ*_*α*_(*x, t*) is the volume fraction of cell type *α* ∈ {*pq, h, e, t, s, p*}, at position *x* and time *t*, 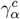 is the consumption rate of the cell type *α*, and *γ*_*α*_ is the production rate of the cell type *α*. In the analysis that follows, we assume that all tumor cells produce VEGF equal to the value calibrated from the cell culture and VEGF expression computational-experimental scenario (see sections 2.4.1 and 2.4.2) but stop producing VEGF whenever they are within the threshold distance from the sprouting vessel. For the calibration of sprout elongation rate in the global analysis, we use a simplified model to calculate the length of the angiogenic sprout to compare with the experimental data and assume the growth of the sprout is independent of VEGF concentration. This simplified model is identical to the full model but with a significantly reduced domain and only one tip cell that grows perpendicular to the parent vessel, allowing removal of the VEGF field, thereby dramatically reducing computational cost while maintaining the angiogenic growth dynamics.

#### 2.3.4 Bayesian Calibration

Bayesian inference provides a statistical framework for calibrating model parameters by accounting for both uncertainties in the mathematical model and the experimental data. Thus, it is an excellent methodology for determining uncertainties in model predictions of experimental outcomes [58–61]. We now summarize the salient features of Bayesian parameter calibration.

Beginning with Bayes’ Rule,

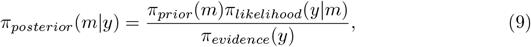

where *m* denotes the model parameters, *y* is the experimental data, *π*_*posterior*_(*m*|*y*) is the distribution of model parameters given the experimental data, *π*_*prior*_(*m*) is the prior distribution of the model parameters, *π*_*likelihood*_(*y*|*m*) is the conditional probability of the experimental data given the model parameters, and *π*_*evidence*_(*y*) is a normalizing factor. These probability density functions (PDFs) provide a holistic description of the system (and therefore) the parameter uncertainty. Since *π*_*evidence*_(*y*) is a normalizing factor, we can rewrite the posterior distribution as,

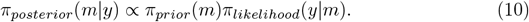

The prior information of the parameters, *π*_*prior*_(*m*), is chosen on a problem specific basis. If only the bounds *a* and *b* of the parameter are known, then the prior is selected as a uniform distribution, *π*_*prior*_(*m*_*i*_) ∼ *U* (*a, b*). Since only the bounds are known in this work, we exclusively use uniform priors. This means that *π*_*prior*_(*m*_*i*_) is constant and we can further simplify Bayes’ Rule to

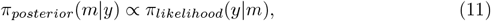

Assuming independent and identically distributed data,

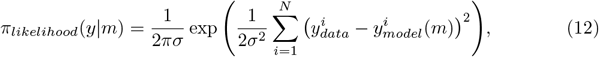

where *σ* is the standard deviation, *N* is the total (combination of spatial and temporal) number of data points, 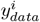 is the experimental data, and 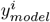 is the model prediction given parameters *m*. This can also be written as,

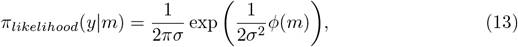

where *ϕ*(*m*) is the misfit to the data. With a given data misfit, Eq. 13 is straightforward to calculate. The posterior distribution for the parameters can be learned by utilizing sampling or quadrature methods to compute *π*_*likelihood*_(*y*|*m*) at specific values of the parameters. Therefore, this framework allows us to use experimental data to learn the updated distribution of parameters. These parameter distributions can then be used to propagate uncertainties in the model predictions for a robust description of prediction uncertainty.

### 2.4 Model calibration and prediction scenarios

We now define the parameter estimation problems in the ODE models (i.e., Eqs. 1 – 4) and the hybrid multiscale model (i.e., Eqs. 5 – 8); the parameters to be calibrated are listed in Table 1. The calibration scenarios results and analyzes are provided at https://github.com/CalebPhillips5/Vascular_calibration.

**Table 1.** Model parameters to be calibrated.

| Parameter | Interpretation | Source |
| --- | --- | --- |
| $\lambda_e^g$ | Growth rate of TIME endothelial cells | Scenario 1 |
| $\theta_e$ | Carrying capacity of TIME endothelial cells | Scenario 1 |
| $\lambda_T^g$ | Growth rate of IBC3 tumor cells | Scenario 1 |
| $\lambda_T^p$ | VEGF production rate by IBC3 tumor cells | Scenario 2 |
| $\lambda_e^c$ | VEGF consumption rate by TIME endothelial cells | Scenario 2 |
| $\theta_v$ | Carrying capacity of VEGF | Scenario 2 |
| $d_{SC}$ | Angiogenic sprout elongation rate | Scenario 3 |
| $f_v$ | VEGF force coefficient | Scenario 3 |
| $d_{tip}$ | Distance between new tip cells | Scenario 4 |
| $d_{sc}^*$ | Local sprout elongation rate | Scenario 5 |

#### 2.4.1 Scenario 1: Calibration of tumor and endothelial cell growth

First, we utilize the hemocytometry and ELISA data to calibrate the parameters in both the ODE systems (Eqs. 1 and 4). We assume endothelial cell growth is independent of the VEGF concentration and can therefore separate the coupled systems into two subsequent calibrations. The parameters calibrated in this scenario are the growth rate of tumor and endothelial cells, and the carrying capacity of endothelial cells. We then define the parameter estimation problem for parameters *m* in both models as: Find *m* such that

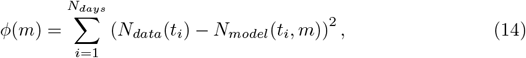

is minimized, *N*_*data*_(*t*_*i*_) and *N*_*model*_(*t*_*i*_, *m*), denote the measured and model prediction of tumor and endothelial cells respectively, *N*_*days*_ as the number of days the VEGF concentration was measured at time points *t*_*i*_. We refer to *ϕ*(*m*) as the data misfit. Since the cell number is dependent on the experimental setup, we use the model and calibrated parameters only to calculate the number of cells over time, which is an input function required to infer the VEGF production and consumption rates for tumor and endothelial cells, respectively, in the coupled ODE systems.

#### 2.4.2 Scenario 2: Calibration and prediction of VEGF concentration

For the VEGF calibration problem, the parameters calibrated in this scenario are the production and consumption rates of VEGF, as well as the carrying capacity of VEGF in the dish (Eqs. 2 and 4). We then define the parameter estimation problem in both models as: Find *m* such that

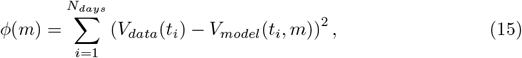

is minimized, where *V*_*data*_(*t*_*i*_) and *V*_*model*_(*t*_*i*_, *m*), denote the measured and model prediction of concentration of VEGF as and respectively, and *N*_*days*_ as the number of days the VEGF concentration was measured. In this scenario, we employ a naïve Bayesian quadrature approach to minimize *ϕ*(*m*) with respect to the parameters *m*. Since the VEGF production and consumption rates are assumed to transcend the experimental scenario (as characteristics of the cells as opposed to characteristics of the environment), we will sample from these distributions moving forward in both the prediction of the VEGF concentration as well as the prediction of the vascular ABM (section 2.4.4. To easily generate samples, we approximate the PDFs as Gaussian distributions.

#### 2.4.3 Scenario 3: Calibration and prediction of global vessel length

We utilize confocal microscopy images to measure the length of angiogenic sprouts over time. This provides a 1D length measurement that can be used to calibrate parameters that govern the growth and development of the sprouts. We selected vessels that can be observed on day 3 and subsequently tracked during the remainder of the experiment to ensure consistent (i.e., the growth of the same sprouts) sprout growth. This is done via a reference mark on the microfluidic platform to align the slices across time. The key parameters are the VEGF force coefficient, *f*_*v*_, and the angiogenic sprout elongation rate, *d*_*sc*_. We then define the parameter estimation problem as: Find m such that

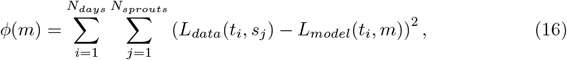

is minimized, where *L*_*data*_(*t*_*i*_, *s*_*j*_) and *L*_*model*_(*t*_*i*_, *m*) are the observed and predicted length of sprout *s*_*j*_ at time point *t*_*i*_, with parameters m at time point *t*_*i*_. *N*_*sprouts*_ was chosen such that the same sprouts were tracked throughout the duration of the experiments. The calibration is solved using a Metropolis-Hastings algorithm available in the QUESO (Quantification of Uncertainty for Estimation, Simulation, and Optimization) library [62]. The prediction of *L*_*data*_(*t*_*i*_, *s*_*j*_) is solved by running the forward model with parameters selected from specific samples used in the Metropolis-Hastings algorithm for calibration and calculating the mean and standard deviation of the resulting quantities of interest, *L*_*model*_(*t*_*i*_, *m*).

#### 2.4.4 Scenario 4: Calibration and prediction of vascular density

To conclude our global analysis, we calculate the vascular density moving away from the parent vessel across time. This provides a global description of the total vascular sprouting to infer the probability distribution function of the distance between new tip cells. We then define the parameter estimation problem as: Find *m* such that

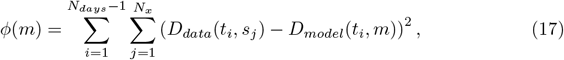

is minimized, where *D*_*data*_(*t*_*i*_, *s*_*j*_) and *D*_*model*_(*t*_*i*_, *m*) are the experimentally estimated (from the confocal microscopy data) and model prediction of the vascular density, respectively, at time *t*_*i*_ and voxel *s*_*j*_. Due to the parent vessel in the microfluidic platform is not perfectly straight and some endothelial cells may migrate short distances from the parent vessel without forming mature vessels, we only utilize voxels *s*_*j*_ that are beyond 15 voxels (*>*35 µm) away from the parent vessel. This calibration scenario is solved by using a naïve quadrature approach as the parameter space is only one dimensional and the computational complexity of the model is prohibitive for sampling. A quadrature approach allows us to utilize parallel resources to run thousands of simulations concurrently, while sampling requires several thousand model runs in serial. For the calibration, we use the day 3, 5, and 7 data and the calibrated probability distribution for distance between new tip cells to predict the vascular density at day 9.

We assess the predictive capabilities of the hybrid multiscale model of tumor angiogenesis at the global and local scales by comparing the model prediction, using calibrated parameter distributions, to the observed data at the final time point in each experiment. The uncertainty in the predictions of vascular density and vascular volume fraction is quantified globally in two cases: the 1-parameter case, using only the parameter distribution for distance between new tip cells (*d*_*tip*_; values sampled from a Gaussian distribution calibrated in Scenario 4), and the 4-parameter case, using the parameter distributions for VEGF production and consumption (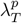 and 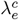), sprout elongation rate (*d*_*sc*_), and the distance between new tip cells (*d*_*tip*_; values are sampled from Gaussian distributions calibrated from Scenarios 2-4). This is accomplished by sampling from the Gaussian fits of the parameter distributions and running the model with the sampled model parameters to calculate the quantities of interest. In the global analysis, we predict the vascular density and vascular volume fraction, while in the local analysis we predict the centerlines of the angiogenic sprouts.

#### 2.4.5 Scenario 5: Calibration of local sprout elongation rate to predict vessel morphology

We employ the confocal microscopy images (see 2.2.3 Image acquisition and processing) to calibrate the hybrid multiscale model to local vascular structures over time. To capture the spatial differences between the model and the data, the optimization problem involves comparing the average distance between the centerlines of the vessels observed in the data and computed by the model. Due to the fact that the local analysis focuses on specific vascular sprouts, we initialize the mathematical model from the data at the first time point (day 3 of the experiment), calibrate the growth of the stalk cells from days 5 and 7, and predict the vascular structure at day 9. We then define the parameter estimation problem as: Find m such that

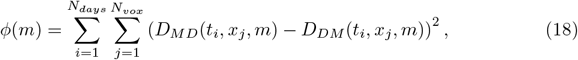

is minimized, where *D*_*MD*_(*t*_*i*_, *x*_*j*_, *m*) is the average distance between the voxels on the centerline of the model to the data and *D*_*DM*_ (*t*_*i*_, *x*_*j*_, *m*) is the average distance between the voxels on the centerline of the data to the model. Across days 5-7 and days 7-9, only two regions had a Dice score above 0.5 and a percentage of overlap above the average of all regions analyzed.

To assess the model prediction of local vessel morphology, we implemented the following steps: 1) take 1000 samples from the Gaussian distribution calibrated from Scenario 5, 2) run the forward model with those 1000 parameter realizations, 3) calculate the centerlines of the forward model solutions, 4) sum over the 1000 centerlines to create a vessel centerline heat map, and 5) threshold the heat map at 10, 50, and 100 to provide the prediction envelope for 1%, 5%, and 10% of the simulations (i.e., voxels covered by 10, 50, and 100 predicted vessel centerlines). The quartiles of the centerline prediction of each region are calculated from MATLAB’s ‘quantiles’.

## 3 Results

### 3.1 Calibration Scenarios 1 and 2: Cell number and VEGF expression

Fig 3 provides an overview of how cell number and ELISA analysis are integrated with the ODE models for the calibration of VEGF production and consumption rates. Cell number was calculated *via* hemocytometer at 24, 48, 72, 120, and 168 hours post seeding. The first time point initialized the TIME and IBC3 growth models and the subsequent four time points were used to calibrate the models for cell number over time (i.e., Eqs. 1 and 3). The number of cells are independent of VEGF concentration allowing the tumor and endothelial cell number calibration to be carried out first with *N*_*e*_(*t*) and *N*_*T*_ (*t*) used as input functions into the production and consumption models of VEGF, respectively. The fit of the growth models is shown in Panels (B) and (C) with growth rates of 8.9 × 10^−3^*s*^−1^ and 3.7 × 10^−2^*s*^−1^ for the IBC3 and TIME cells, respectively, and a carrying capacity of 2.67 × 10^5^ for the TIME cells. In the ELISA experiments, every 24 hours for 7 days, VEGF concentration is measured, and media is replaced; note that this action causes the total concentration of VEGF in the plate to return to baseline (i.e., 1100 pg/mL) after each measurement. The VEGF experiments for TIME and IBC3 cells are carried out in different plates, so the calibration of each cell line is not coupled. We utilize naïve Gaussian quadrature to calculate the probability distribution of VEGF production by TIME cells, as well as the VEGF consumption and carrying capacity of the IBC3 cells. The maximum likelihood values were obtained using MATLAB’s ‘lsqnonlin’ function, and we then utilized a parameter sweep using 100 values for each of the three parameters (total of a million points) taken about the maximum likelihood values. The resulting posterior distributions are shown in Panel (D). The fit Gaussian distributions are given by

**Figure 3.**
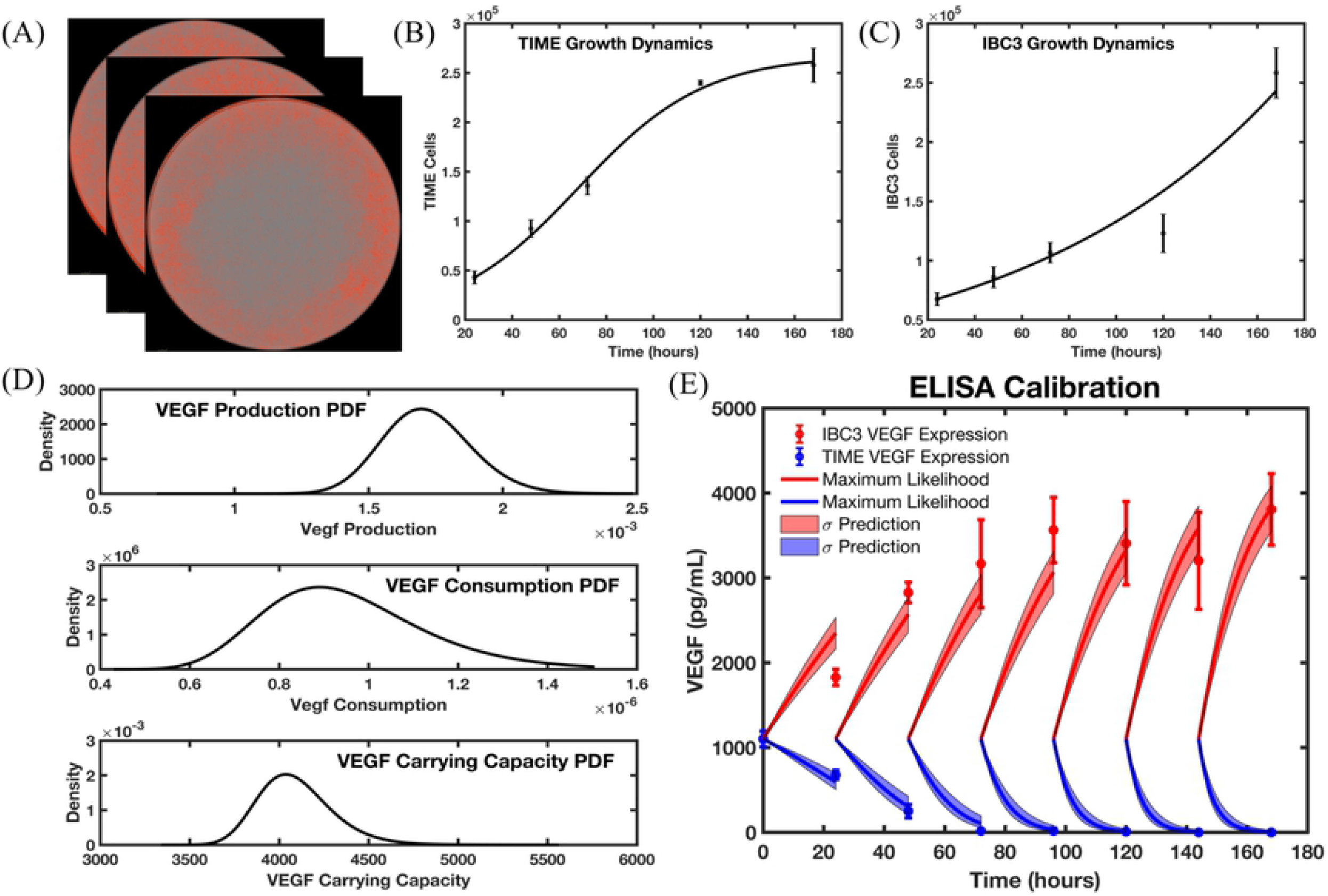
Calibration and prediction scenario for the cell culture and VEGF expression data. Panel (A) shows an example of temporally resolved IncuCyte images used for determining the tumor and endothelial cell number. Panels (B) and (C) show the observed number of TIME and IBC3 cells over time with the calibrated logistic growth model for TIME cells and exponential growth for IBC3 cells. Panel (D) shows the posterior distribution function of parameters inferred using the IncuCyte and ELISA data. The calibrated parameters from Panels (B) and (C) are utilized in the models of VEGF secretion and consumption by IBC3 and TIME cells, respectively, to calibrate VEGF production, consumption, and carrying capacity. These posterior distributions are sampled to propagate uncertainty in the ODE models of VEGF secretion and consumption, with the one standard deviation confidence intervals shown in Panel (E). The VEGF expression models for tumor and endothelial cell have an average relative error of 11% and 16.7%, respectively. The calibrated parameter distributions for VEGF production and consumption are integrated back into the hybrid multiscale angiogenesis model for subsequent analysis.

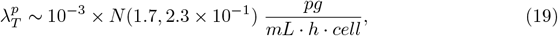

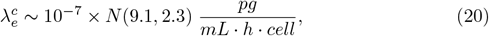

and

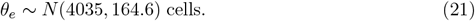

In Panel (E), the IBC3 and TIME VEGF concentrations (with standard deviations) are shown as the red and blue dots, respectively. The maximum likelihood value for each parameter is

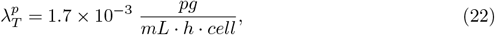

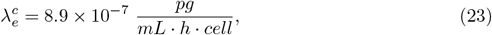

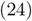

and

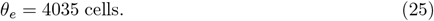

The means (and associated *σ* prediction envelope) of the best fit of TIME and IBC3 VEGF models are shown in blue and red, respectively. The calculated distributions and maximum likelihood values of VEGF production and consumption rates will be used in subsequent calibration and prediction scenarios described below in Sections 3.2-3.4. The relative errors between the models and experimental measurements are summarized in Table 2.

**Table 2.**
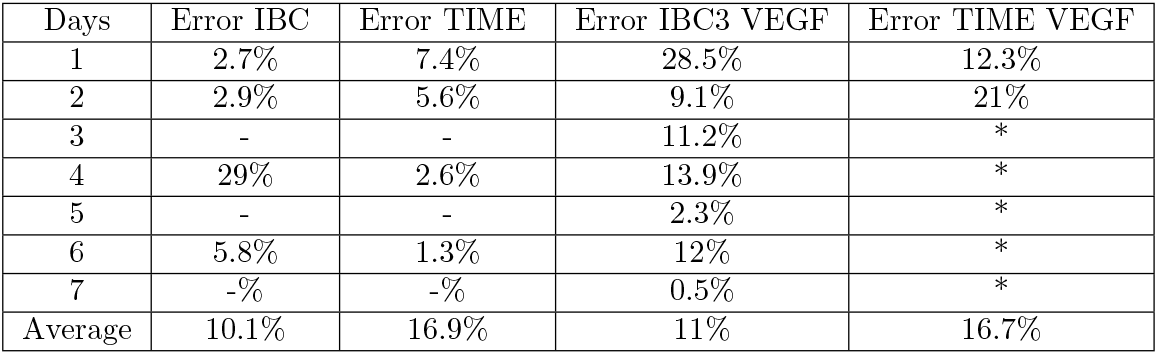
Error between model output and cell culture and VEGF expression data. All errors are relative errors between the model output and experimental data. The dashes (-) denote where the data is not available to calculate the quantity, and the asterisk (*) denotes when the data records a value of less than 2% of the baseline concentration of 1100 pg/mL, thereby precluding computing the relative error (i.e., it would be near infinite).

| Days | Error IBC | Error TIME | Error IBC3 VEGF | Error TIME VEGF |
| --- | --- | --- | --- | --- |
| 1 | 2.7% | 7.4% | 28.5% | 12.3% |
| 2 | 2.9% | 5.6% | 9.1% | 21% |
| 3 | - | - | 11.2% | * |
| 4 | 29% | 2.6% | 13.9% | * |
| 5 | - | - | 2.3% | * |
| 6 | 5.8% | 1.3% | 12% | * |
| 7 | -% | -% | 0.5% | * |
| Average | 10.1% | 16.9% | 11% | 16.7% |

### 3.2 Calibration Scenario 3: Calibration and prediction of global sprout elongation rate

Fig 4 shows the calibration of length measurements of angiogenic sprouts (imaged *via* confocal microscopy) to the sprout lengths predicted by the vascular ABM. Sprout length measurements are taken on days 3, 7, 11, 15, and 19 after the 72-hour flow protocol (see 2.2.2 Vascularized 3D *in vitro* microfluidic platforms). Panel (A) depicts the sprout length measurements at day 11. Panel (B) shows the mean and standard deviation of the data in black and the standard deviation of the prediction of the vascular ABM in light red using the calibrated parameters, along with the mean given by the black line. The posterior distributions of stalk cell divide time, VEGF force, and a hyper parameter standard deviation (which encompasses both the experimental and model uncertainty) are shown in Panels (C), (D), and (E), respectively. The error in the calibration is shown in Table 3.

**Figure 4.**
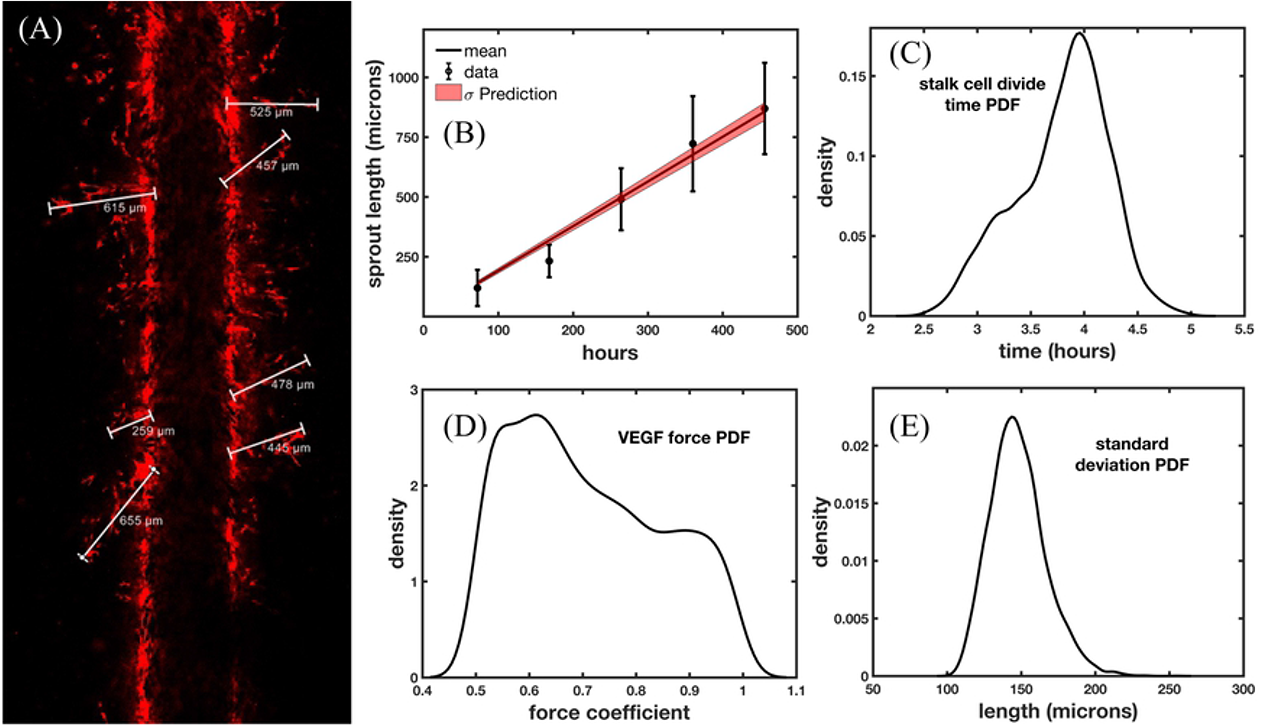
Calibration and prediction scenario for sprout elongation rate and sprout length measurements. Panel (A) displays the Day 11 length measurements from the confocal microscopy images of the angiogenic vasculature in the 3D microfluidic platform outlined in the experimental methods of Scenario 3 (Section 2.2.2. The length measurements over time are depicted in Panel (B), where the dots show the mean of the measurements, and the error bars show the standard deviation. The red region depicts the standard deviation of the prediction of the calibrated mathematical model with calibrated posterior distributions of each parameter shown in Panels (C), (D), and (E). Panel (C) depicts the posterior distribution of the stalk cell divide time, Panel (D) depicts the posterior distribution of the VEGF Force, and Panel (E) depicts the posterior distribution of the standard deviation which has been calibrated as a hyperparameter in this scenario. The average error of predicted sprout length over time is 15.3%. The calibrated stalk cell divide time posterior distribution is utilized in the subsequent global analysis.

**Table 3.** Error in calibration and prediction of sprout length.

| Days | Observed sprout length (microns) | Predicted sprout length (microns) | Predicted error sprout length (%) |
| --- | --- | --- | --- |
| 3 | 119.8±75.8 | 140.2±8.9 | 17.1 |
| 7 | 232.4±67.7 | 318.6±14.6 | 37.1 |
| 11 | 490.6±129.5 | 497.6±29.5 | 14.6% |
| 15 | 722.7±198.7 | 676.0±29.5 | 6.3% |
| 19 | 869.9±191 | 856.8±37.6 | 1.5% |
| Average | - | - | 15.3% |

The maximum likelihood values for stalk cell divide time (*d*_*sc*_), VEGF force coefficient (*f*_*v*_), and standard deviation (*σ*_*s*_) are 3.96 h, 0.61, and 143.5 µm, respectively. The stalk cell divide time and VEGF force values are utilized in the calibration of distance between new tip cells. Approximating these PDFs as Gaussians, the parameters are *d*_*sc*_ ∼ *N* (3.96, 0.44), *f*_*v*_ ∼ *N* (0.71, 0.14), and *σ*_*s*_ ∼ *N* (147.8, 18.6). The Gaussian fit of *d*_*sc*_ is utilized moving forward to assess the uncertainty in the model prediction of vascular density and volume fraction. Going forward, we will ignore the uncertainty contribution of the VEGF force as its primary role is to maintain physiological sprouting that does not break the vessel but balances the forces to allow vessel elongation.

### 3.3 Scenario 4: Calibration and prediction of vascular density

Fig 5 presents the calibration of the distance between new tip cells by utilizing the vascular density of angiogenic sprouts imaged in the microfluidic platform. Panel (A) depicts a representative binary image of the angiogenic sprouts in the microfluidic platform, and Panel (B) shows the calculated density of the angiogenic sprouts. These are aligned together with the ABM so the spatially varying vascular density can be compared between the model and the data. In Panel (C), we show the calibrated PDF of the distance between new tip cells in blue and the Gaussian fit of the PDF, which may be readily sampled for analyzing the uncertainty in the prediction of the model. In Panels (D) - (F), we show days 3, 5, and 7 of the calculated density of the left and right sides of the microfluidic platform (blue and magenta, respectively), and the model best fit obtained during calibration over days 3 and 5 (red). The day 7 model best fit is the temporally evolved solution of the model best fit calibrated from days 3 and 5. The vertical dashed line represents the location from which we begin to make use of the vascular density observed in the microfluidic platform for the calibration. That location is chosen to minimize the effects of migratory endothelial cells close to the channel wall that do not form mature angiogenic sprouts. The Gaussian fit of the PDF is *d*_*tip*_ ∼ *N* (243.9, 18.1) microns. The error in the calibration and prediction is shown in Table 3.

**Figure 5.**
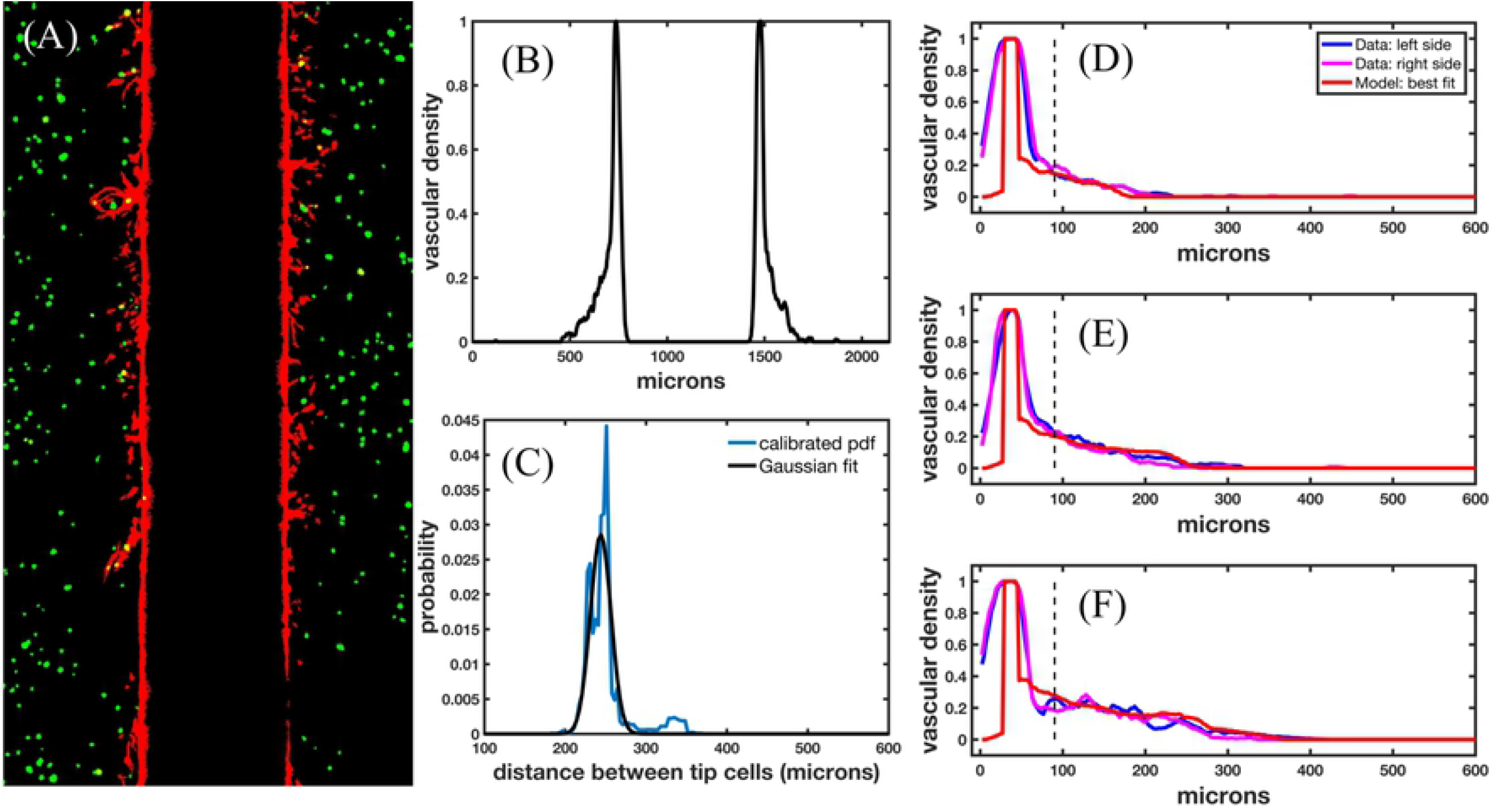
Calibration of distance between tip cells. Panel (A) displays a representative RGB binary image (day 7) from the confocal microfluidic platform used to calculate the vascular density, shown in Panel (B). Panel (C) depicts the calibrated PDF and Gaussian fit of the distance between new tip cells. Panels (D) - (F) show the density calculated from the left and right side of the microfluidic platform shown in blue and magenta, respectively, and the best fit of the model shown in red for days 3, 5, and 7, respectively. The vertical dashed lines represent the location away from the parent vessel that we begin to use for calibration (i.e., we utilize the voxels to the right of the dashed vertical line, see Scenario 4, Section 2.4.4). The relative error of the best fit in the calibration for days 3, 5, and 7 are 23.5%, 11.1%, and 18.5% respectively.

We utilize 1000 samples from the probability distribution of the distance between new tip cells, as well as the probability distributions obtained in scenarios 2 and 3, to rigorously quantify the uncertainty in the predictability of the model. The prediction results are shown in Fig 6. In Panels (A) - (C), we show the 95% prediction confidence intervals propagating uncertainty using one parameter distribution for distance between new tip cells shaded in light red and using four parameter distributions (distance between new tip cells, VEGF production and consumption rates, and stalk cell growth rate) shown in light blue. The vascular density calculated from the left and the right sides of the microfluidic platform are shown in blue and magenta, respectively, and the model best fit is shown in red. The vertical black dotted line denotes the distance away from the vessel that we begin to use for calibration. The total uncertainty (summed over space) was 97.4%, 66.2% and 45.2% higher in the four-parameter case than the 1-parameter case on days 3, 5, and 7, respectively. Panel (D) shows the calculated average vascular fraction from the data in black, and the predicted average vascular fraction for the one parameter (blue) and four parameter (red) uncertainty analyses. In Panel (E), the vascular fraction of the data, 1-parameter, and 4-parameter scenarios are shown in blue, orange, and yellow, respectively.

**Figure 6.**
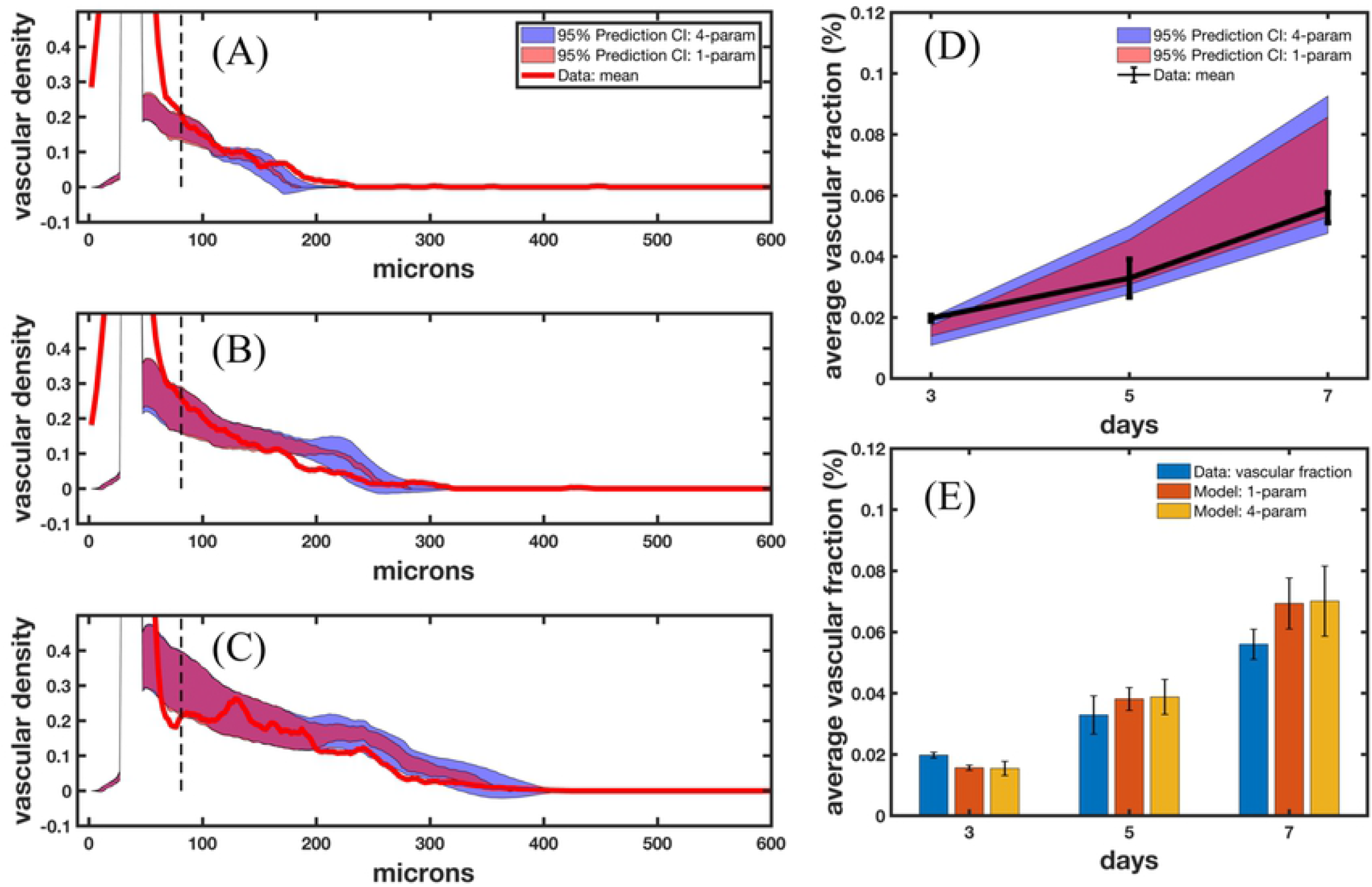
Prediction of vascular density. Panels (A) - (C) show the uncertainty in the model prediction assuming uncertainty in one parameter calibrated from scenario 4 (shaded in light red) and four parameters (shaded in light blue) calibrated from calibration scenarios 2, 3, and 4, along with the mean of the data. In the 1-parameter case, we consider only the distance between new tip cells with a PDF shown in Panel (C) of Fig 5. In the 4-parameter case, we consider distance between new tip cells, VEGF production and consumption rates (shown in Panel (D) of Fig 3), and stalk cell growth rate (shown in Panel (B) of Fig 4). Panel (D) presents the volume fraction calculated from the data in black, and the predicted volume fraction from the 1-parameter and 4-parameter case in light red and blue, respectively. Panel (E) depicts the prediction of volume fraction compared to the data (blue), and the 1-parameter and 4-parameter cases in orange and yellow, respectively, with the corresponding standard deviations shown in black. The average error in vascular volume fraction is 20.2% and 21.7% of the 1-parameter and 4-parameter case, respectively. We also note the drastic increase in uncertainty moving away from the parent vessel in the 4-parameter case (Panel (A) 120-180 microns, Panel (B) 200-280 microns, and Panel (C) 200-400 microns), highlighting the effects of the VEGF production and consumption rates and the sprout elongation rate on the uncertainty in the prediction.

### 3.4 Scenario 5: Calibration of local sprout elongation rate to predict vessel morphology

While sprout length, vascular density, and volume fraction over time are reasonable metrics for analyzing global features of vascular structure, additional local analysis is needed to assess the model’s ability to recapitulate local features in the vascular network. Fig 7 shows the local region selected from the microfluidic platform for the calibration of stalk cell growth rate. Confocal images are taken on days 3, 5, 7, and 9, and processed in MATLAB using a Dice score and area threshold to select regions for local calibration (see 2.2.3 Image acquisition and processing). Once selected by our sorting algorithm, we calculate the skeletonization of the local vascular structures using a Zhang-Suen parallel thinning algorithm [ref]. We use the day 3 skeletonization, shown in Panel (A) of Fig 8 to initialize the model (Panel (D)) by segmenting the tumor cells from the surrounding region and calculating the trajectories of the tip cells shown in the data to formulate the initial conditions of the ABM that match the skeletonization of the data. The stalk cell growth rate is calibrated by running the model forward and comparing the centerline distance between the data at days 5 and 7 (Panels (B) and (C), respectively) and the vasculature predicted by the model with the maximum likelihood stalk cell growth rate, shown in Panels (C) and (F). The resulting probability distribution function of stalk cell growth rate (blue) and the Gaussian fit (black) are shown in Panel (G). The maximum likelihood value for stalk cell growth rate in this scenario is 10.86 hours and the Gaussian fit is *d*_*sc*_ ∼*N* (10.85, 2.4) hours. We then take 1000 samples from this Gaussian distribution and calculate the resulting vasculature for days 5, 7, and 9 as shown in Fig 8.

**Figure 7.**
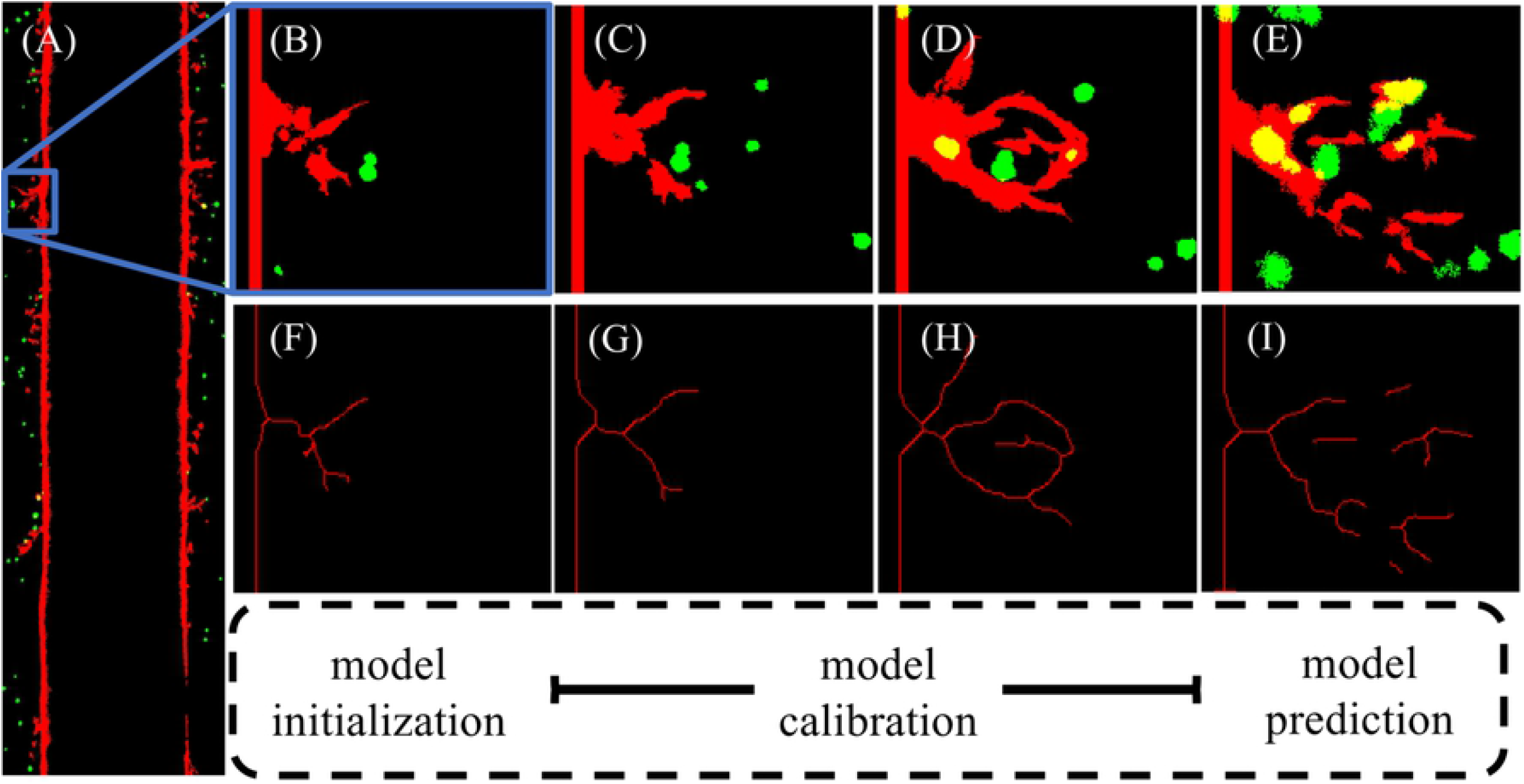
Vascular regions for local calibration. Panel (A) shows a RBG binarized image of vascular structure of day 3 in the microfluidic platform. Panels (B) - (E) depict a specific local region segmented over time (blue box in Panel (A)) with vessels shown in red and tumor cells shown in green from days 3, 5, 7, and 9. Panels (F) - (I) show the corresponding centerline segmentation of Panels (B) – (C), computed from a parallel thinning algorithm. The local analysis flowchart is shown by model initialization using the day 3 centerline, model calibration utilizing days 5 and 7, and model prediction of the centerline from day 9.

**Figure 8.**
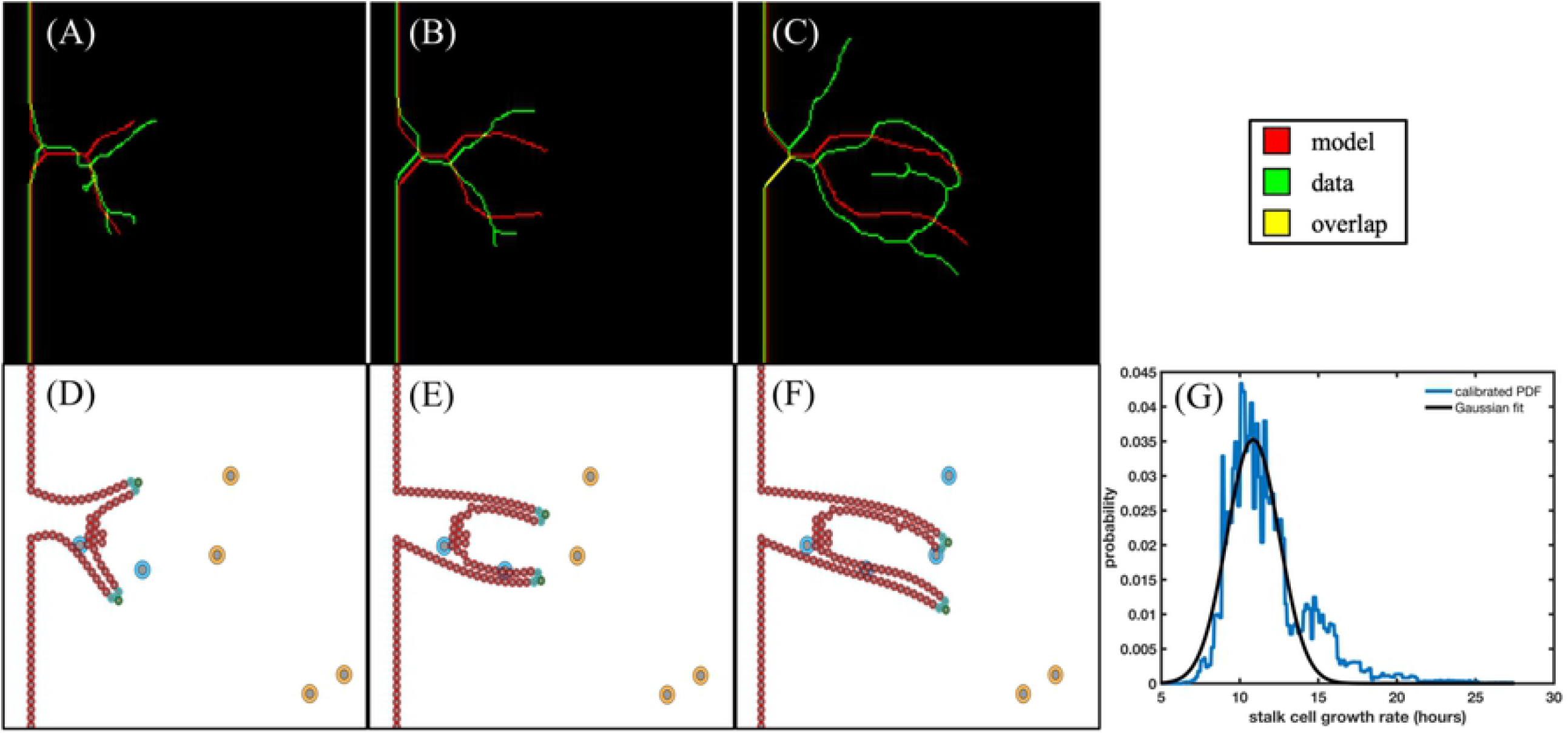
Local region 1: calibration of stalk cell growth rate. Panels (A) - (C) show the centerlines calculated from the model best fit (red), the data (green), and the overlap (yellow) on day 3 (used to inform the initial conditions), day 5, and day 7 (used to calibrate the stalk cell growth rate). The ABM is shown in Panels (D)-(F) with tip cells in green, stalk cells in cyan, endothelial cells in red, tumor cells releasing VEGF in orange, and tumor cells not releasing VEGF in blue. In Panel (G), we show the calibrated PDF and the Gaussian fit of the stalk cell growth rate. The best fit model recapitulates the general structural features of the data without allowing additional sprouts to form. We also note that from day 7 (depicted in Panel (F)) to day 9, only the two tumor cells in the bottom right of the domain continue to release VEGF, guiding the sprout migration to the bottom right of the domain.

In Fig 9, the columns denote day 5, 7, and 9, respectively. The rows depict the 1%, 5%, and 10% prediction envelope (top to bottom, described in 2.2.3 Image acquisition and processing). In Panel (B), the data and the model calculated centerlines form an anastomosis, predicted in 3.3% of simulations. Anastomosis is also predicted on day 9 in Panels (C), (F), and (I) in 16.7% of simulations, recapitulating the vessel structure observed in the data at day 7. In Panel (J), the prediction of the average centerline distance from model to data is calculated from the 1000 samples of the distribution in Fig 8 Panel (G). Panel (K) presents the normalized average centerline distance from the data to model (normalized to the length of the sprout in day 3).

**Figure 9.**
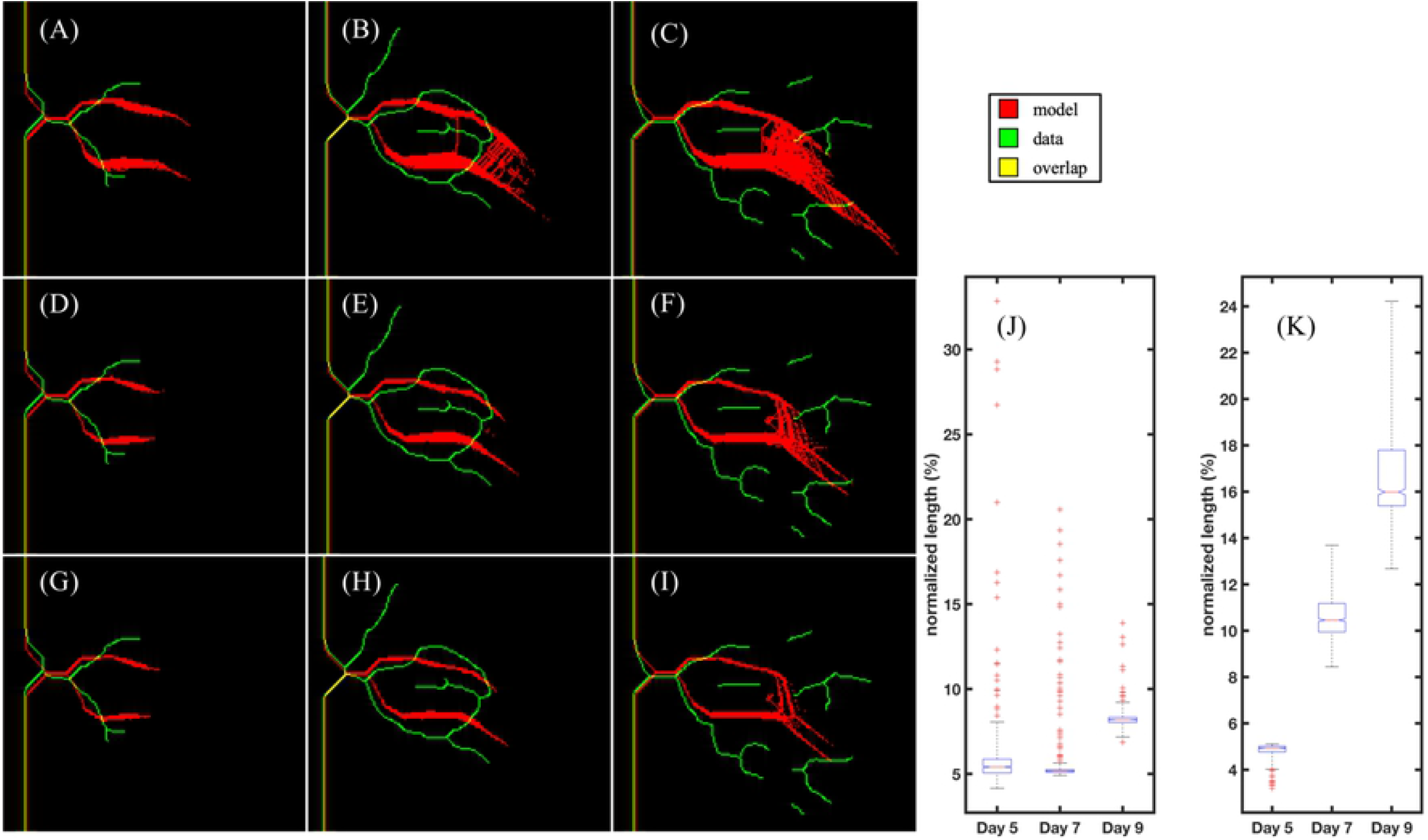
Local region 1: prediction. Each column depicts the data (green), model prediction (red), and overlap of the two (yellow) on days 5, 7, and 9, respectively. Each row shows the predicted centerlines using three different thresholds: 99% prediction (top row), 95% prediction (middle row), and 90% prediction (bottom row) of the simulations. While the data exhibits vessel anastomosis at Day 7, the model prediction only depicts angiogenesis in the 1% prediction at Day 7 (Panel (B)) and Day 9 (Panels (C), (F), and (I)). Panels (J) and (K) shows the prediction of average centerline value from model to data and from data to model, respectively. The average centerline distance is normalized by the length of the longest sprout in this region at day 3. This results in a normalized length of less than 10% from model to data and a normalized length of less than 20% from data to model. However, at day 9 the complexity in the vascular network, specifically the vascular remodeling that eliminates the anastomosis, is beyond the capabilities of the model to predict.

In Figs 10 and 11, we show the results of the calibration and prediction, respectively, of local region 2. In Fig 10, Panels (A) - (C) show the best fit (red), the centerlines of the data (green), and the overlap (yellow), with the corresponding hybrid ABM prediction of days 3, 5, and 7 shown in Panels (D) - (F), respectively. Panel (G) shows the calibrated PDF (black) and the Gaussian fit of the local stalk cell growth rate (blue). The maximum likelihood value is 72.1 hours and the Gaussian fit is *d*_*sc*_ ∼ *N* (78.86, 10.83) hours. In Fig 11, we take 1000 samples from the Gaussian fit of stalk cell divide time and calculate the 1%, 5%, and 10% prediction envelopes of the model, shown in Panels (A)-(C), (D)-(F), and (G)-(I), respectively. The prediction quartiles of both local regions are shown in Table 5.

**Figure 10.**
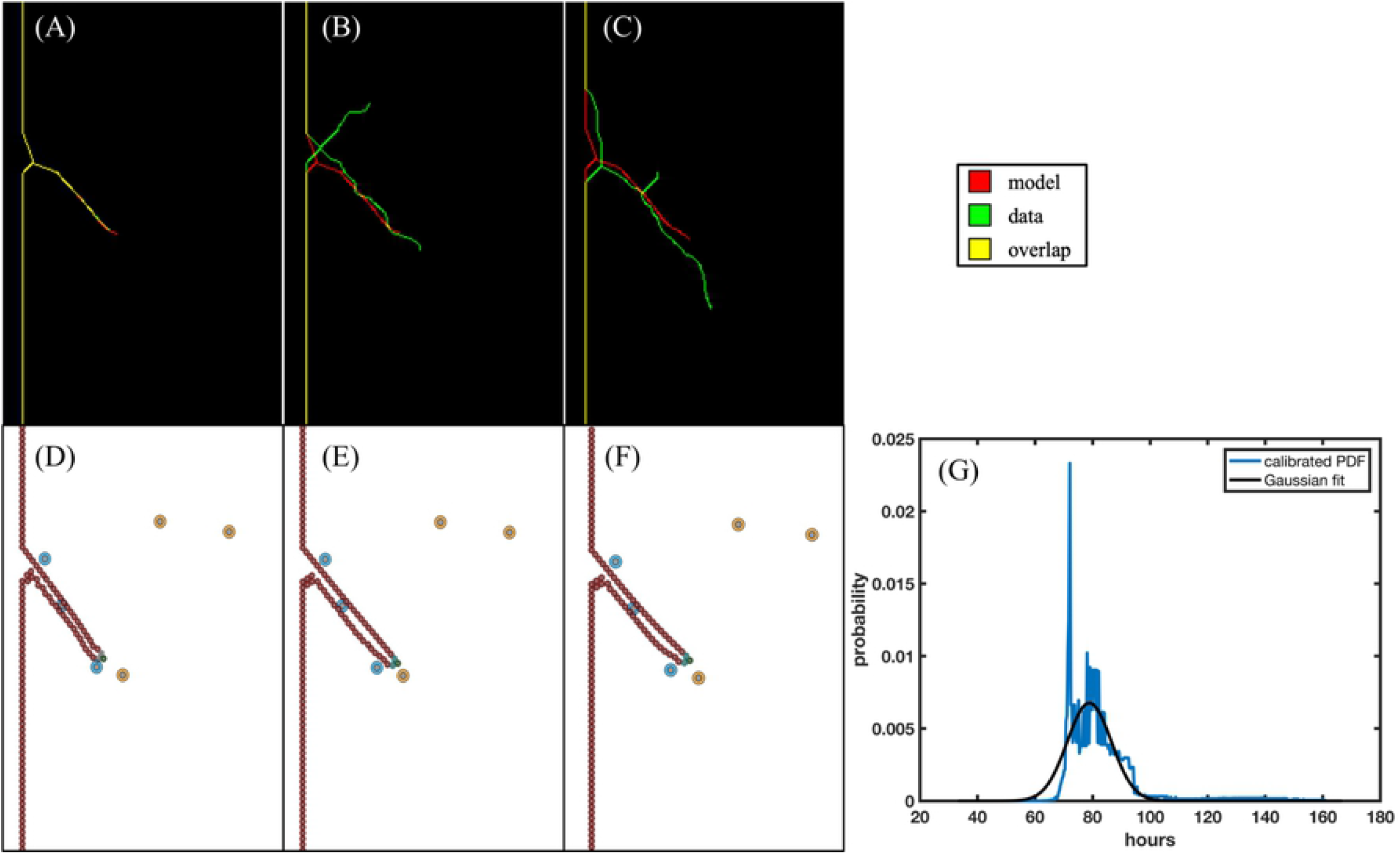
Local region 2: calibration of stalk cell growth rate. Panels (A) - (C) show the centerlines calculated from the model best fit in red, the data in green, and the overlap in yellow of day 3 (used to inform the initial conditions), Day 5, and day 7 (both used to calibrate the stalk cell growth rate). The agent-based model is shown in Panels (D)-(F) with tip cells in green, stalk cells in cyan, endothelial cells in red, tumor cells releasing VEGF in orange, and tumor cells not releasing VEGF in blue. In Panel (G), we show the calibrated PDF and the Gaussian fit of the stalk cell growth rate. The mean of the local stalk cell divide time (∼ 80 hours) is significantly higher than the global stalk cell divide time calibrated at ∼4 hours. This highlights the effect of the hypoxic tumor cells in the top right portion of the domain, as a higher calibrated stalk cell divide time would cause growth toward this region of space, while the data continues toward the bottom right of the domain.

**Figure 11.**
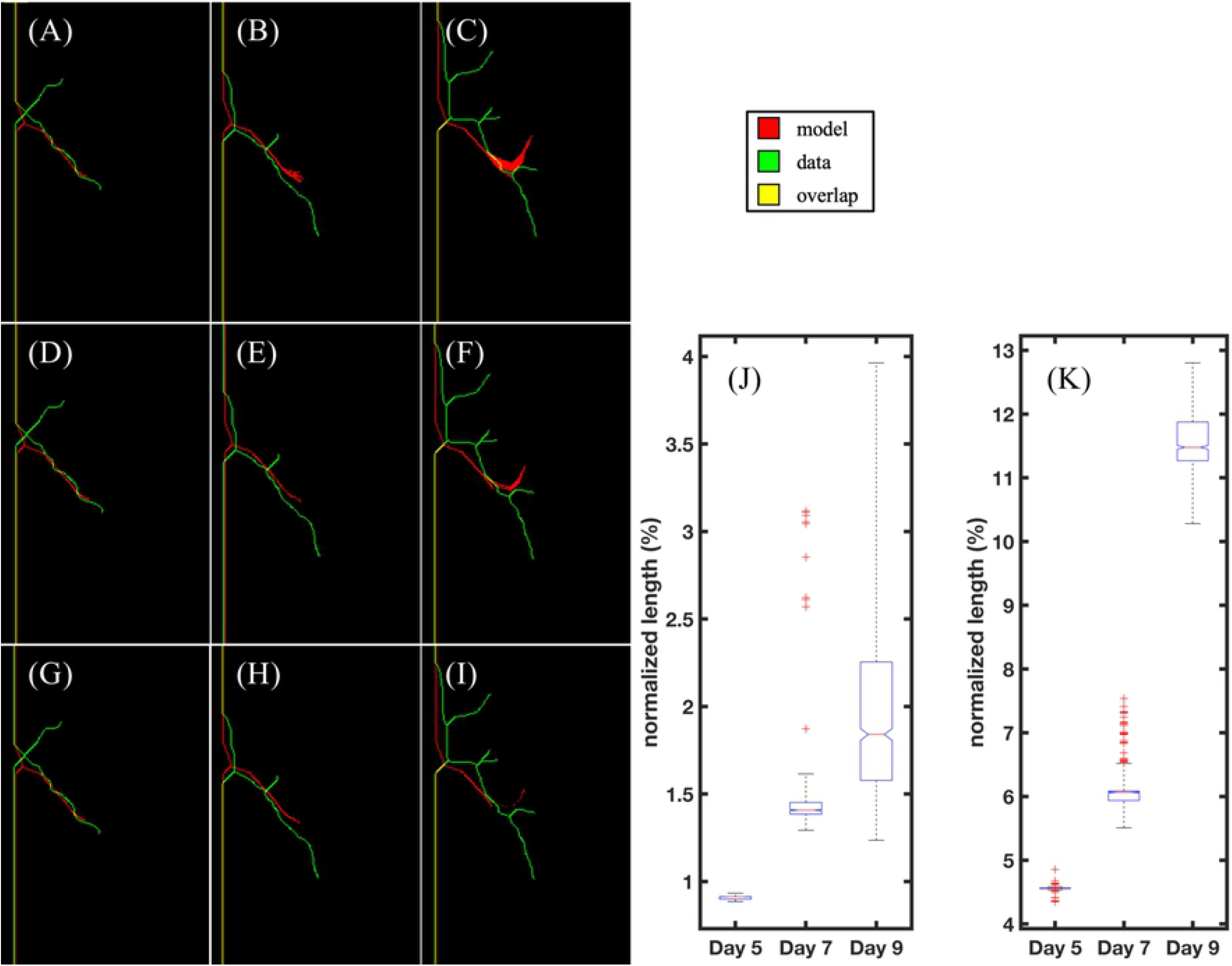
Local region 2: prediction. Each column depicts the data (green) and model prediction (red), with overlap in yellow, of days 5, 7, and 9, respectively. Each row shows the voxels predicted by the centerlines using different thresholds, with the voxels in the 99% prediction (top row), 95% prediction (middle row), and 90% prediction (bottom row) of the simulations. In Panel (C), the direction of vessel growth in the model shows the VEGF gradient going away from the direction of data and toward the hypoxic cells shown in Fig 9 Panel (F). This leads to the overall directionality of the vessel to be misaligned with the data. Panels (J) and (K) shows the prediction of average centerline value from model to data and from data to model with an averaged normalized length difference of less than 2% and 12%, respectively. The average centerline distance is normalized by the length of the longest sprout in this region at day 3.

## 4 Discussion

We have rigorously calibrated a hybrid, multiscale model of tumor angiogenesis, both with global and local vascular structure, with experimental data obtained from an *in vitro* vascularized tumor platform and tested the ability of the model to make accurate longitudinal predictions. This was done by first utilizing hemocytometry and ELISA to calibrate mathematical models of cell growth and VEGF expression, respectively, yielding parameter distributions for VEGF production and consumption rates (Fig 3). The maximum likelihood values are used in subsequent calibration and prediction scenarios. Secondly, we analyzed global features such as sprout length, vascular density, and vascular volume fraction over time, to calibrate the global growth rate of angiogenic sprouts and the distance between new tip cells, shown in Figs 4 and 5. We then reported the errors and rigorously accounted for the uncertainty in model predictions, in Table 4 and Fig 6, respectively. The mean error in the prediction of the 1- and 4-parameter cases was 21% and did not increase with time. Finally, we analyzed the ability of the model to recapitulate local vascular structure by segmenting and skeletonizing specific vessels over time, initializing the model with confocal images at day 3, calibrating with images at day 5 and 7, and testing the predictability of the model against day 9 observations, as summarized by Fig 7. Our calibrated hybrid multiscale model, informed by hemocytometry and VEGF expression data, can recapitulate local vascular structure longitudinally in our *in vitro* microfluidic platform. The calibration and prediction error, shown in Table 5, was on the order of 10% normalized centerline distance. The normalized length of the centerline distance from data to model increases across time and indicates that the hybrid ABM using the initial conditions of two tip cells (see Fig 9 (D)), while quantitatively and qualitatively matching the description of vasculature at day 7, fails to capture the complexity observed at day 9. This is partially due to the restriction of new tip cells in the local calibrations, as we do not allow for new sprouts. However, the complexity across time points of the local sprouts dramatically increases and requires further study from both a modeling and experimental standpoint.

**Table 4.** Error in calibration and prediction of vascular density. We compare the relative error in the calibration best fit, the 1-parameter, and 4-parameter prediction cases.

| Days | Calibration best fit % error | 1-Parameter prediction mean % error | 4-Parameter prediction mean % error |
| --- | --- | --- | --- |
| 3 | 23.6 | 20.9 | 21.9 |
| 5 | 11.1 | 15.9 | 18 |
| 7 | 18.5 | 23.8 | 25.2 |
| Average | 17.7 | 20.2 | 21.7 |

**Table 5.** Error in calibration and prediction of vascular density. We compare the relative error in the calibration best fit, the 1-parameter, and 4-parameter prediction cases.

| Region | Quartiles | $C_{dist}(m, d)$ | | | $C_{dist}(d, m)$ | | |
| --- | --- | --- | --- | --- | --- | --- | --- |
|  |  | 25% | 50% | 75% | 25% | 50% | 75% |
| 1 | Day 5 | 5.1% | 5.4% | 5.9% | 4.8% | 4.9% | 5.0% |
|  | Day 7 | 5.1% | 5.2% | 5.3% | 10% | 10.5% | 11.2% |
|  | Day 9 | 8.0% | 8.2% | 8.3% | 15.4% | 16% | 17.8% |
| 2 | Day 5 | 0.7% | 0.7% | 0.7% | 3.6% | 3.6% | 3.6% |
|  | Day 7 | 1.1% | 1.1% | 1.1% | 4.7% | 4.8% | 4.8% |
|  | Day 9 | 1.2% | 1.5% | 1.8% | 8.9% | 9.0% | 9.3% |

To the best of our knowledge, this work represents the first study to rigorously calibrate an agent-based model of tumor angiogenesis with multimodal experimental data to forecast vascular growth that is then directly compared to the corresponding data. Importantly, other efforts have pioneered the integration of data in other ways [63, 64]. For example, Perfahl *et al*. employed multiphoton microscopy to image angiogenic vasculature in an *in vivo* dorsal skin fold chamber [7]. These images furnished the initial conditions of a cellular automaton model of vascular tumor growth, with blood flow, and vascular remodeling. However, no model calibration or comparison to experiments was performed. Similarly, in Xu *et al*., photoacoustic imaging of a murine xenograft model initialized a phase-field model of tumor angiogenesis [38]. While the authors did not utilize time-resolved vascular data, this experimental-computational approach showcases a first step toward longitudinally integrating *in vivo* data into angiogenesis models, through initialization, calibration, and ultimately, prediction. The present study incorporates initialization, calibration, and prediction through utilizing *in vitro* data; however, the experimental and computational approaches in this contribution can be made model-agnostic (and, even, data-agnostic). In particular, all the calibration methods were based on gradient free methods, and all the solvers are independent of the model, data, and the system under investigation.

A key experimental limitation of the present study is the lack of longitudinal measures of various proteins (e.g., VEGF) and cell markers (e.g., to assign endothelial cells as tip, stalk, phalanx phenotype) in the *in vitro* microfluidic platform. We have addressed this by devising smaller calibration and prediction scenarios (with corresponding experiments) that can isolate salient phenomena (e.g., VEGF production from tumor cells and growth of stalk cells) that we assume are independent of the domain. Another area for advancement is to determine the generalizability of both the local and global calibration results by applying the approach to multiple microfluidic platforms. We utilized one microfluidic platform over time, yielding two regions for global analysis and two local regions for calibration and prediction. We note that in our global analysis in Scenario 4, as we utilize an ABM for angiogenesis, the model cannot be uniquely initialized from day 3 or day 5 and is thus restarted from day 0 with no angiogenic vasculature for each simulation. This is because the model is not continuous and cannot be populated from the vascular density at days 3 or 5, but the endothelial cells in the ABM must start from a parent vessel. A key limitation in the modeling is that our calibration was only performed on 2D data. As out of plane effects could play a significant role in vessel sprouting, future efforts will be focused on extending the formalism to 3D. The main limitation to extending to 3D is the characterization of the angiogenic sprouts by endothelial cells that make up both sides (in 2D) of the sprout. This can be overcome by describing the vessels as nodes (which describe the radius of the vessel) connected by edges (which describe the directionality of the vessels), an approach taken by [65, 66]. In our future work, we aim to utilize more experimental replicates to further calibrate and validate our model and to devise experimental protocols with anti-angiogenic and radiation therapies to model the effects of these treatments on angiogenic sprouts *in vitro* [67].

## 5 Conclusion

We have calibrated and determined the ability of an agent-based model to make accurate predictions of tumor angiogenesis by systematically incorporating data of different scales to inform model parameters. To the best of our knowledge, this represents the first effort to calibrate a mechanism-based mathematical model to spatially and temporally-resolved experimental data of angiogenesis, thereby enabling predictions of future vessel development that could then be directly tested against observation.

## Acknowledgments

We thank the National Cancer Institute for funding *via* U01CA174706, R01CA186193, U01CA253540, and U24 CA226110. We also thank the Cancer Prevention and Research Institute of Texas (CPRIT) for funding *via* RR160005. T.E.Y. is a CPRIT Scholar in Cancer Research. M.N.R acknowledges support from the VPR Research and Creative Grant and the Walker Department of Mechanical Engineering at The University of Texas at Austin. We also thank the Texas Advanced Computing center for providing high-performance computing resources.

